# Brain disconnectivity mapping of post-stroke fatigue

**DOI:** 10.1101/2020.11.13.380972

**Authors:** Kristine M. Ulrichsen, Knut K. Kolskår, Geneviève Richard, Dag Alnæs, Erlend S. Dørum, Anne-Marthe Sanders, Sveinung Tornås, Jennifer Monereo Sánchez, Andreas Engvig, Hege Ihle Hansen, Michel Thiebaut de Schotten, Jan E. Nordvik, Lars T. Westlye

**Affiliations:** NORMENT, Division of Mental Health and Addiction, Oslo University Hospital & Institute of Clinical Medicine, University of Oslo, Norway; Department of Psychology, University of Oslo, Norway; Sunnaas Rehabilitation Hospital HT, Nesodden, Norway; Bjørknes College, Oslo, Norway; Department of Neurology, Oslo University Hospital, Norway; Brain Connectivity and Behaviour Laboratory, Sorbonne Universities, Paris, France; Groupe d’Imagerie Neurofonctionnelle, Institut des Maladies Neurodégénératives-UMR 5293, CNRS, CEA University of Bordeaux, Bordeaux, France; CatoSenteret Rehabilitation Center, Son, Norway; Faculty of Health, Medicine and Life Sciences, Maastricht University, Netherlands; Department of Radiology and Nuclear Medicine, Maastricht University Medical Center, Netherlands; Department of Nephrology, Oslo University Hospital, Ullevål, Norway; KG Jebsen Centre for Neurodevelopmental Disorders, University of Oslo, Norway

**Keywords:** Stroke, post-stroke fatigue, MRI, brain mapping, disconnectome, lesion

## Abstract

Stroke patients commonly suffer from post stroke fatigue (PSF). Despite a general consensus that brain perturbations constitute a precipitating event in the multifactorial etiology of PSF, the specific predictive value of conventional lesion characteristics such as size and localization remain unclear. The current study represents a novel approach to assess the neural correlates of PSF in chronic stroke patients. While previous research has focused primarily on lesion location or size, with mixed or inconclusive results, we targeted the extended structural network implicated by the lesion, and evaluated the added explanatory value of a disconnectivity approach with regards to the brain correlates of PSF. To this end, we estimated individual brain disconnectome maps in 84 stroke survivors in the chronic phase (≥ 3 months post stroke) using information about lesion location and normative white matter pathways obtained from 170 healthy individuals. PSF was measured by the Fatigue Severity Scale (FSS). Voxel wise analyses using non-parametric permutation-based inference were conducted on disconnectome maps to estimate regional effects of disconnectivity. Associations between PSF and global disconnectivity and clinical lesion characteristics were tested by linear models, and we estimated Bayes factor to quantify the evidence for the null and alternative hypotheses, respectively. The results revealed no significant associations between PSF and disconnectome measures or lesion characteristics, with moderate evidence in favor of the null hypothesis. These results suggest that symptoms of post-stroke fatigue are not simply explained by lesion characteristics or brain disconnectome measures in stroke patients in a chronic phase, and are discussed in light of methodological considerations.

## Introduction

Between 25 and 85 percent of stroke survivors experience post stroke fatigue (PSF) (Cumming, Packer, Kramer, & English, 2016), described as an excessive and debilitative tiredness that can be unrelated to strain and not ameliorated by rest (UK Stroke Association, 2020; de Groot, Phillips, & Eskes, 2003). Persistent PSF can be highly distressing, negatively impacting quality of life (de Bruijn et al., 2015; Naess, Waje-Andreassen, Thomassen, Nyland, & Myhr, 2006) and preventing social participation and attendance to rehabilitation programs (Nadarajah & Goh, 2015). PSF is associated with both poor functional outcome and increased mortality (Glader, Stegmayr, & Asplund, 2002), and a recent meta-analysis revealed that the prevalence increases with time since stroke (Cumming et al., 2018). Early detection, prevention and treatment of fatigue might thus have positive effects on the overall outcome of stroke rehabilitation and quality of life. As such, identification of risk factors is important to facilitate detection and individual tailoring of rehabilitation programs.

A number of biological, psychological, demographical and social risk factors for PSF have been suggested. Briefly; reduced physical function, female sex, depression (Lerdal et al., 2011; Aarnes, Stubberud, & Lerdal, 2019), pain, sleep disturbances (Naess, Lunde, Brogger, & Waje-Andreassen, 2012) and certain lesion characteristics (Mutai, Furukawa, Houri, Suzuki, & Hanihara, 2017; Wai Kwong Tang et al., 2014; Wai Kwong Tang et al., 2010) are among the identified risk factors. Depression is the most consistently correlated factor (Ponchel, Bombois, Bordet, & Hénon, 2015; Wu, Barugh, Macleod, & Mead, 2014), and, although PSF is generally conceptualized as an independent condition, the nature of the relationship between fatigue and depression has been debated. Although efforts have been made to disentangle the two (Douven et al., 2017; Høgestøl et al., 2019; Kunze, Zierath, Drogomiretskiy, & Becker, 2014), the clinical overlap is substantial (Cumming et al., 2018). The use of advanced brain imaging to detect the brain correlates of the two clinical syndromes may facilitate our understanding of the phenomena through identification of both common and specific brain mechanisms (Høgestøl et al., 2019).

Despite a general consensus that the lesion and the associated brain perturbations following the stroke constitute causal factors for PSF, little is known about the predictive value of key lesion characteristics such as extent and neuroanatomical distribution. Fatigue is more prevalent following a minor stroke compared to a transient ischemic attack (TIA) (Naess et al., 2012; Winward, Sackley, Metha, & Rothwell, 2009), suggesting that the vascular lesion itself is of importance with regards to fatigue. Further, stroke survivors describe the fatigue experienced after stroke as qualitatively different than fatigue before stroke or normal tiredness (Thomas, Gamlin, De Simoni, Mullis, & Mant, 2019). Lastly, fatigue is a common sequela or symptom in several neurological conditions, i.e. traumatic brain injury, multiple sclerosis and postpoliomyelitis, jointly referred to as “central fatigue” (Chaudhuri & Behan, 2000, 2004).

Studies examining associations between lesion characteristics and fatigue in stroke survivors have generated mixed findings. In line with the hypothesis of fatigue caused by nervous system disruptions (Chaudhuri & Behan, 2000, 2004), basal ganglia infarcts have been identified as predictors of fatigue (Wai Kwong Tang et al., 2010) and caudate infarcts were more frequent in patients with than without PSF (W. K. Tang et al., 2013). Further, infratentorial infarcts have been associated with increased risk of fatigue (Snaphaan, Van der Werf, & de Leeuw, 2011), as have right hemisphere lesions, and brainstem and thalamic lesions (Mutai et al., 2017). Yet the relationship between fatigue and lesion location remains unresolved (De Doncker, Dantzer, Ormstad, & Kuppuswamy, 2018), and several studies report no consistent associations between lesion characteristics and fatigue (Choi-Kwon, Han, Kwon, & Kim, 2005; Ingles, Eskes, & Phillips, 1999; Mead et al., 2011).

It is conceivable that clinical symptoms following a stroke are not mediated primarily by the localization of the lesion, but rather by the functional neuroanatomy of the extended brain networks that are affected by the lesion and degree of preserved network function (Bartolomeo & de Schotten, 2016; Nordin et al., 2016). Neuroimaging suggests that many psychiatric and neurologic symptoms are related to complex brain networks of anatomically distant but connected regions (Fox, 2018) that are vulnerable to injuries in a range of locations. Through processes like diaschisis (remote neurophysiological changes or dysfunctions of a distant region caused by a focal injury (Carrera & Tononi, 2014; von Monakow, 1914)), disconnection (Geschwind, 1974) and transneuronal degeneration (Cowan, 1970), stroke lesions may affect brain function and behavior in ways not readily predicted by the location or size of the damaged tissue. For example, functional MRI has revealed that functional network disturbances are observed between remotely connected cortical areas in both the unaffected and affected hemisphere (Rehme & Grefkes, 2013), and abrupted connectivity may cause impairments that are functionally similar to direct tissue necrosis (Bonilha et al., 2014). Probing the extended brain network characteristics involved in a lesion and its associations with outcome may therefore provide theoretically and clinically relevant information of the functional neuroanatomy of specific symptoms post stroke and other brain disorders.

Recent large-scale collaborative neuroimaging efforts have resulted in remarkable advances in the characterization of the human brain “connectome” and “disconnectome” (Thiebaut de Schotten, Foulon, & Nachev, 2020), providing highly valuable roadmaps for studies attempting to link symptoms, lesions and brain networks. Various approaches for lesion network mapping (Foulon et al., 2018; Fox, 2018) are now being applied to neurological conditions such as mania symptoms (Cotovio et al., 2020), amnesia (Ferguson et al., 2019), and Alzheimer’s disease (Darby, Joutsa, & Fox, 2019). In the context of stroke lesions and fatigue, targeting network disconnections in addition to lesion characteristics may advance our understanding on the relationship between brain perturbations and fatigue beyond what is revealed by traditional lesion-symptom mapping.

To evaluate the added explanatory value of a disconnectivity based approach with regards to the brain correlates of PSF, we quantified lesion disconnectivity indirectly using information about normative white matter pathways in the healthy population to estimate individual structural disconnection (disconnectome) maps in 84 stroke survivors in the chronic phase. The maps were created by a tractography-based procedure (Foulon et al., 2018) yielding voxel-wise probability of structural disconnection of white matter tracts (Salvalaggio, De Filippo De Grazia, Zorzi, Thiebaut de Schotten, & Corbetta, 2020).

Associations between disconnectome maps and PSF (assessed by the Fatigue Severity Scale (FSS)), were examined using permutation testing. Due to the substantial overlap and interaction between fatigue and depression and the possibility of common mechanisms across these conditions, all voxel-wise analyses were done with a) fatigue scores, b) depression scores (measured using the Pittsburg Health Questionnaire (PHQ-9) (Spitzer, Kroenke, Williams, & Patient Health Questionnaire Primary Care Study, 1999) and c) fatigue and depression scores combined. The above described analyses were repeated on the binarized lesion maps, reflecting a traditional voxel-based lesion symptom mapping (VSLM) approach (Bates et al., 2003). In addition, we estimated the global disconnectivity for each patient, and tested for correlations with FSS, using Bayes factor to quantify evidence for the null hypothesis. Finally, in agreement with a more traditional, clinical approach, we applied linear models to test for associations between PSF and stroke related factors such as stroke location (right hemisphere, left hemisphere, brainstem, cerebellum, or both hemispheres), months since stroke, stroke severity (using National Institute of Health Stroke Scale (NIHSS; Lyden et al., 2009) score at discharge as a proxy for clinical severity) and etiology as defined by the stroke subtype classification system Trial of Org 10172 in Acute Stroke Treatment (TOAST; Adams Jr et al., 1993).

Due to a lack of previous studies applying a disconnectivity approach to PSF, we remained agnostic about the specific brain networks involved and performed a whole-brain analysis. Based on recent work demonstrating the benefits of targeting network projections of a lesion (Griffis, Metcalf, Corbetta, & Shulman, 2019; Thiebaut de Schotten et al., 2020), and the notion that many psychiatric and neurological conditions correspond more closely to brain networks than specific regions (Fox, 2018), we expected the disconnectivity based approach to demonstrate higher sensitivity to PSF than conventional lesion-related approaches.

## Materials and methods

### Study participants

Table 1 summarizes demographic and clinical information. We recruited 84 stroke patients from the Geriatric Department, Diakonhjemmet Hospital, the Stroke Unit, Oslo University Hospital and Bærum Hospital. A subsample of the patients (n=66) participated in a longitudinal intervention study examining the effects of cognitive training and tDCS on cognitive function (see Kolskaar et al. (2020) for details). All data reported in the current study were collected prior to the intervention. Criteria were ischemic or hemorrhagic stroke in a chronic phase defined as ≥3 months since admission, age above 18 years, no MRI contraindications and no other severe neurological conditions.

**Table 1.**

| *Sample characteristics* | Patients (n=84) |  | Control group (n=155) |  |  |  |
| --- | --- | --- | --- | --- | --- | --- |
| *Demographic and clinical information* | Mean(SD) | Min-Max | Mean(SD) | Min-Max | t (p) | BF** |
| Age | 65.8(12.6) | 24 - 87 | 64.7(12.3) | 24 - 92 | 1.0 (.279) | 0.16 |
| Males/females (count) | 60/24 |  | 111/44 |  |  |  |
| Education in years | 14.5(3.4) | 7 - 30 | 15.7 (3.3) | 6 - 25 | 0.6 (.513) | 0.6 |
| FSS | 3.9(1.5) | 1 - 7 | 2.9 (1.3) | 1 - 6 | -3.9 (<.001) | 1911 |
| PHQ-9 | 5.0(4.5) | 0 - 21 | 3.2 (3.0) | 0 - 15 | -3.2 (.001) | 30 |
| Montreal cognitive assessment (MoCA) | 26.3(2.4) | 19 - 30 | 27.4 (1.7) | 22 - 30 | 3.0 (.002) | 171 |
| *Stroke related patient information* |  |  |  |  |  |  |
| NIHSS at hospital discharge | 1.1 (1.2) | 0 - 6 |  |  |  |  |
| TOAST classification for ischemic stroke* | Large artery atherosclerosis (26)<br>Small vessel occlusion (26)<br>Cardioembolism (13)<br>Other determined etiology (6)<br>Undetermined etiology (13) |  |  |  |  |  |
| Lesion location | Brainstem/cerebellum (17)<br>Left Hemisphere (26)<br>Right Hemisphere (34)<br>Both Hemispheres (6) |  |  |  |  |  |
| Months since stroke | 22.0 (11.9) | 3 - 45 |  |  |  |  |

\*all but one patient suffered ischemic stroke
\*\*BF = Bayes factor.
Both frequentist and Bayesian statistics are reported, in line with current pragmatic recommendations (Keyesers, Gazzola, & Wagenmakers, 2020).

Healthy individuals >18 years were recruited through newspaper ads, word-of-mouth and social media (Dørum et al., 2020; Richard et al., 2018). Exclusion criteria included a history of stroke, neurological or psychiatric disease, medications with significant effects on central nervous system function and MR contraindications.

Healthy controls and stroke patients were matched on age and sex, using *MatchIt* in R (Stuart, King, Imai, & Ho, 2011) and the default method *nearest*. Applying a ratio of 2:1 (two controls selected for each patient), healthy participants were collected from a pool of 341 controls (age 24 – 92), resulting in an age- and sex matched control group of 155 individuals (mean age = 64.7, SD = 12.3, 44 females).

All participants were screened using Montreal cognitive assessment (MoCA). A score below 25 may indicate mild cognitive impairment (Nasreddine, 2020; Nasreddine et al., 2005). One included patient had a MoCA score of 19 and one healthy control a score of 22, but further neuropsychological assessments indicated sufficient cognitive function and did not reveal contraindications for study participation.

All participants provided informed consent prior to enrollment. The study was approved by the Regional Committee for Medical and Health Research Ethics, South-East Norway.

### Fatigue Severity Scale (FSS)

Subjective fatigue was measured by the FSS (Krupp, LaRocca, Muir-Nash, & Steinberg, 1989), which is a self-report scale consisting of 9 statements about impact of fatigue on daily life. Degree of agreement is indicated on a seven-point Likert scale (lowest possible total score 7, highest score 63). FSS is one of the most frequently used instruments for measuring fatigue in neurological conditions (Cumming et al., 2016) and has demonstrated reasonable psychometric qualities (Whitehead, 2009). A commonly adapted threshold for clinical fatigue is a mean score of >= 4 (total score >= 36) (Krupp et al., 1989; Nadarajah & Goh, 2015; Schepers, Visser-Meily, Ketelaar, & Lindeman, 2006), where a higher score is suggested to indicate a moderate to high impact of fatigue (Schepers et al., 2006).

### Patient Health Questionnaire (PHQ-9)

Depression symptoms were measured by the depression module of the PHQ-9, in which occurrence of depressive symptoms corresponding to the DSM-IV criteria is rated on a 9-item Likert scale. Scores range from 0 (not at all) to 3 (nearly every day), yielding a minimum score of zero and a maximum score of 27. A cutoff score of ≥ 10 has demonstrated acceptable sensitivity and specificity for depression (Kroenke, Spitzer, & Williams, 2001).

To establish the degree of symptom load in the stroke sample compared to healthy individuals, we conducted two tailed t-tests on group differences in FSS and PHQ-9.

### MRI acquisition

Patients were scanned at Oslo University Hospital on a 3T GE 750 Discovery MRI scanner with a 32-channel head coil. We collected structural (T1w, FLAIR), functional (resting-state and task-based fMRI) and diffusion MRI data. For lesion demarcation used in the present analysis T1-weighted images were collected using a 3D IR-prepared FSPGR (BRAVO) sequence (TR: 8.16 ms; TE: 3.18 ms; TI: 450 ms; FA: 12°; voxel size: 1×1×1 mm; slices: 188; FOV: 256 & 256, 188 sagittal slices), and T2-FLAIR with the following parameters: TR: 8000 ms; TE: 127 ms, TI: 2240; voxel size: 1×1×1 mm).

### Lesion demarcation

Lesions were demarcated in native space, using the Clusterize toolbox (de Haan et al., 2015) with SPM8 running under Matlab R2013b (The Mathworks, Inc., Natick, MA). Lesions were traced by trained personnel (a physician and a radiographer), based on hyperintensities and visible damage on FLAIR images, and guided by independent neuroradiological descriptions of dMRI/FLAIR images (see Dørum et al. (2020) for details). The binarized lesion masks were registered to MNI space using nearest neighbor interpolation, using the transformation parameters obtained using the T1w data. To register the FLAIR images to the T1 images, we applied a linear transformation with 6 degrees of freedom. T1 images were registered to MNI152 space by linear affine transformation with 12 degrees of freedom. Figure 1 displays a probabilistic map of lesion overlap across patients.

**Figure 1.**
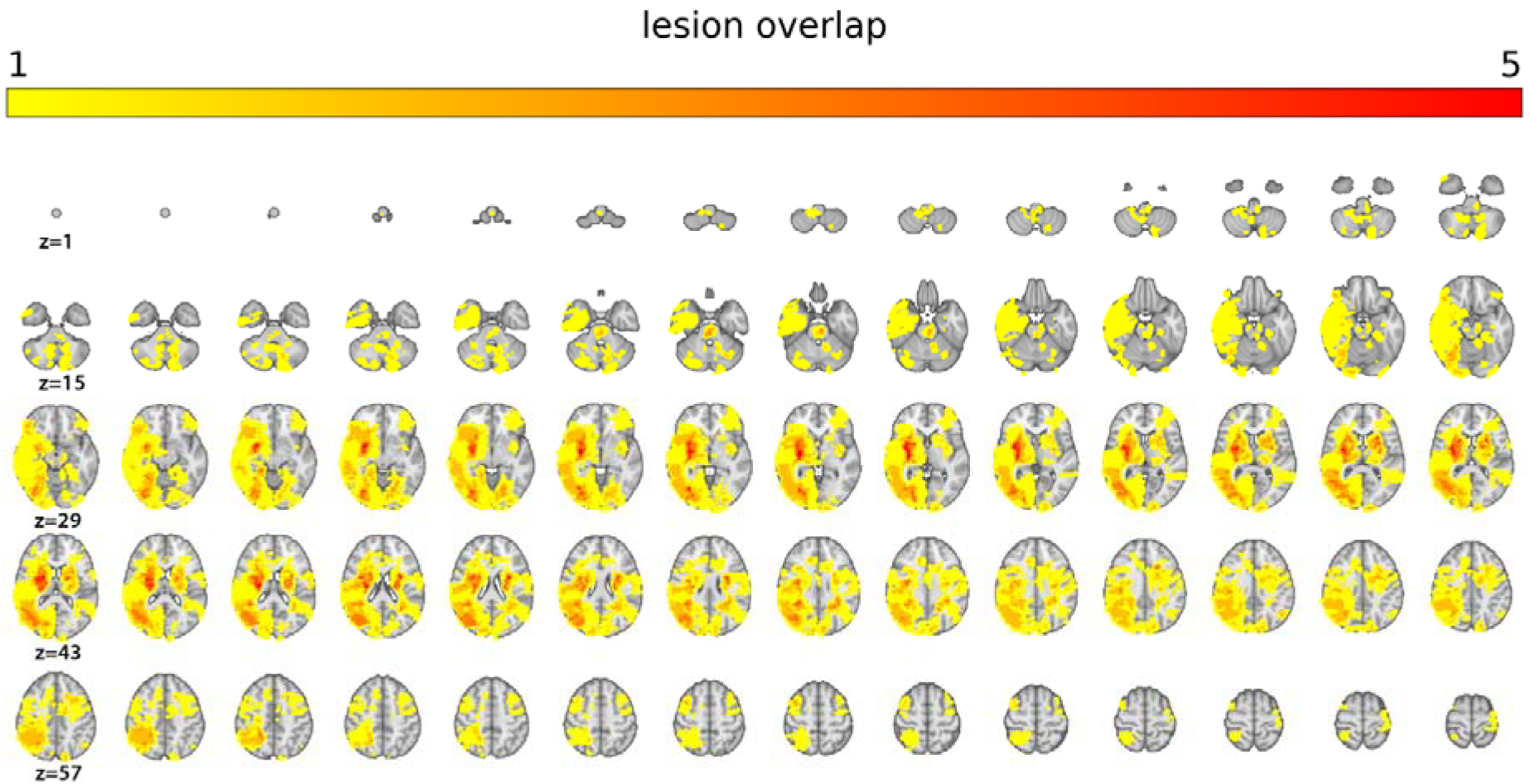
Heatmap displaying lesion overlap across stroke patients by 70 slices (2 mm thickness) from z(voxel)=1 to z=70. Maximal overlap was 8, but for illustration purposes, the color scale saturates at 5.

### Disconnectome maps

To calculate the disconnectome maps we used an automated tractography-based procedure (Foulon et al., 2018) implemented in the *BCBtoolkit disconnectome maps* (“Brain Connectivity Behaviour Toolkit (BCBtoolkit),”). Briefly, a training set based on full-brain tractography data obtained from a normative group of 170 individuals from the Human Connectome Project 7T data (HCP 7T) was used to track fibers passing through each lesion. Using affine and diffeomorphic deformations (Avants et al., 2011; Klein et al., 2009), each patient’s MNI 152 space lesions were registered to each control’s native space, and used as seed for the tractography in Trackvis (Wang, Benner, Sorensen, & Wedeen). Subsequently, the tractography was transformed to visitation maps, binarized and registered to MNI152 space, before a percentage overlap map was produced by summarizing each point in the normalized healthy subject visitation maps. The resulting disconnectome maps indicate a voxel-wise probability of lesion-related disconnection ranging from 0 -100 %. We computed two simple summary measures of disconnection severity, defined for each patient as a) mean voxel intensity across the individual disconnectome map and b) number of voxels within the individual disconnectome map with intensity >.5 (reflecting 50% probability of disconnection).

### Statistical analysis

Voxel wise analyses on disconnectome maps and binarized lesions were done by non-parametric permutation-based inference as implemented in the FSL randomise tool (Winkler, Ridgway, Webster, Smith, & Nichols, 2014). Within the framework of the general linear model (GLM), linear effects of fatigue and depression (indicated by total score on FSS and PHQ, respectively) were tested in separate models, covarying for age and sex. A supplementary FSS-model controlled for depression by excluding patients scoring above clinical threshold on PHQ (remaining n = 74). To comply with a more common clinical definition of PSF, we re-ran the model on dichotomized fatigue variables defined as either a) a mean FSS score of ≥ 4, consistent with the common cut off value, or b) the upper tertile of FSS total score (contrasted with the lowest tertile), reflecting the possibility that more extreme scores demonstrate increased sensitivity to fatigue related brain correlates. One additional model tested the effect of fatigue and depression combined, applying the total of the z-normalized sum scores (zPHQ + zFSS) as predictor. For each contrast, 5000 permutations were performed. Results were thresholded by threshold free cluster enhancement (TFCE, Smith and Nichols (2009)) and considered significant at p<.05, two tailed, corrected for multiple comparisons using permutation testing. One patient suffered a very large stroke and constituted an outlier in terms of number of affected voxels (~8 SDs above the mean). Main analyses were therefore repeated with this patient excluded.

Subsequent statistical analyses were performed using R version 3.4.0 (R Core Team, 2017). In a follow-up analysis aiming to increase sensitivity to clinical measures and evaluate the relationship between global disconnectivity and fatigue, we computed two disconnection severity measures, defined for each patient as a) mean voxel intensity across the individual disconnectome map and b) number of voxels within the individual disconnectome map with intensity >.5 (reflecting 50% probability), and correlated these with FSS and PHQ-9. To quantify the evidence in favor of the null and alternative hypothesis, we applied Bayes factor hypothesis testing, in line with current recommendations (Keysers et al., 2020). We applied the BayesFactor package (Morey, Rouder, Jamil, & Morey, 2015) with default priors. For transparency, key analyses were run with different priors.

To test for associations with clinical, stroke-related characteristics (TOAST classification, months since stroke, lesion volume and lesion location), we applied linear models with FSS score as dependent variable, controlling for age and depressive symptoms. Lesion location was clustered by four categories – right or left hemisphere, both hemispheres or brainstem/cerebellum. Stroke variables were added subsequently, allowing for model comparison by Bayes factor for each added variable. We applied the lmBF function from the BayesFactor package to compute Bayes factors. As an additional test of the added predictive value of global disconnectivity measures compared to clinical stroke characteristics, we also estimated the models with mean voxel intensity across the individual disconnectome map and number of voxels within the individual disconnectome map with intensity >.5.

## Results

### Fatigue and depression in the stroke sample compared to healthy controls

Figure 2 shows the distributions of FSS and PHQ score in each group. 48 percent of the stroke patients reported clinically significant fatigue (mean FSS > 4), compared to 23 percent of the control participants. Severe fatigue (mean FSS > 6) was reported by 9 percent of the patients and 1 percent of the healthy controls. A two-tailed, two sample t-test (ttestBF in BayesFactor, with default Cauchy prior) provided compelling evidence (Bayes Factor: BF) > 150) for higher total FSS scores in the patient group (mean = 35) compared to healthy controls (mean = 26, median posterior *δ* = −8.5, 95% credible interval (CI)= [−11 – −5]), relative to the null hypothesis. Stroke patients (mean = 5.0) also reported higher levels of depression symptoms on the PHQ than controls (mean = 3.2). 18 percent of the patients scored 10 or higher, indicating clinical depression, compared to 2.4 percent of the controls. The corresponding Bayes factor provided strong evidence for a group difference in PHQ sum score (BF = 20, 95% CI = [−2.5 – −0.5]).

**Figure 2.**
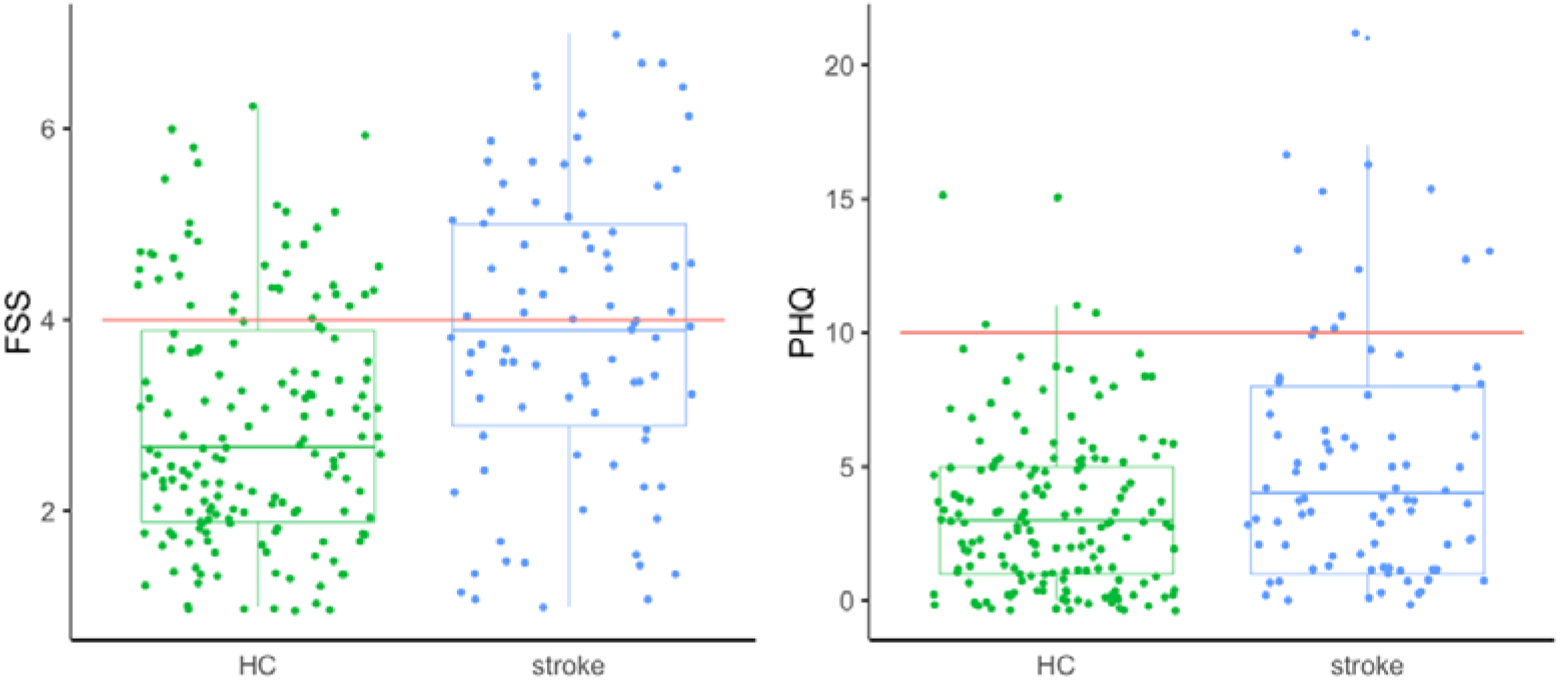
Distributions and group differences in FSS and PHQ for healthy controls (HC) and stroke patients. Red line denotes cut off value for clinically significant symptom load.

Among patients, Bayes factor estimation for linear correlations provided strong evidence (BF > 150) for a positive association between FSS and PHQ-9, (median posterior *δ* = 0.67, 95% CI = [0.54 – 0.58]), suggesting more depressive symptoms with increasing fatigue. There was only anecdotal support for an association between FSS and age (BF = 1.75, median posterior *δ* = 0.21, 95% CI = [0.00 – 1.42]).

### Permutation based analyses on disconnectivity maps and lesions

Figure 3 shows a selection of stroke lesions and the associated disconnectome maps, for illustrative purposes.

**Figure 3.**
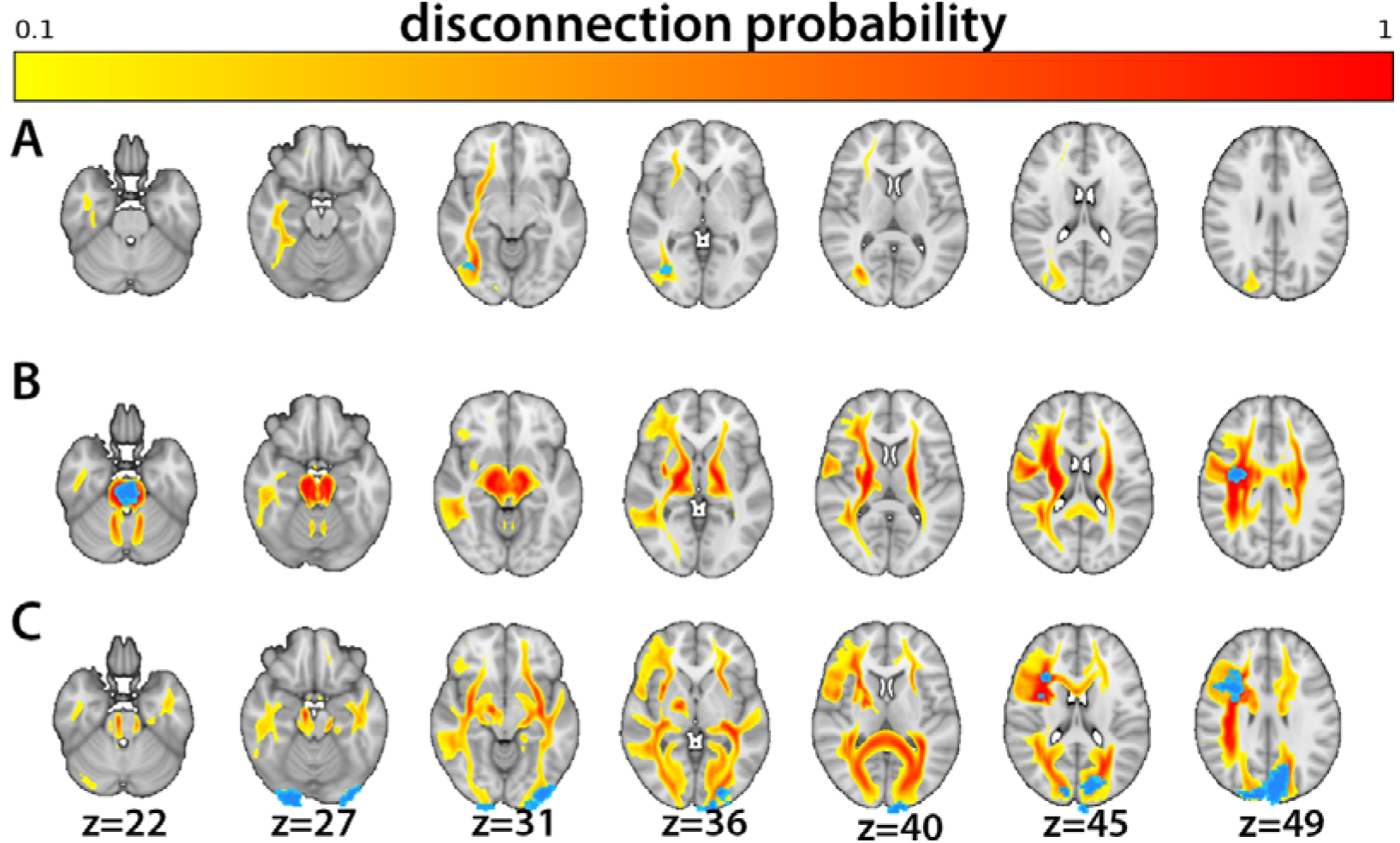
Individual lesions (blue) and associated disconnectome maps (yellow-red). Probability for disconnection ranges from 10 (yellow) to 100 (red). Patient **A**: right cerebral white matter lesion, Patient **B:** brain stem lesion, Patient **C:** left and right cerebral cortex and white matter lesions.

Permutation testing revealed no significant associations between the disconnectivity maps and the clinical measures (FSS, PHQ-9, FSS/PHQ combined), or of fatigue status defined by either a) mean FSS score of ≥ 4, or b) by the lowest vs highest FSS tertile. Results were mirrored in the permutation tests on binarized lesion maps, revealing no association between the clinical measures (fatigue, depression or fatigue/depression combined, or on group defined by fatigue status (mean FSS score ≥ 4). Due to the considerable reduction in sample size when including only the lowest and highest FSS tertile (n = 56), we did not repeat this analysis on the binarized lesion maps.

### Associations between global summary measures of disconnectivity and clinical measures

Both measures of global disconnectivity (mean value in disconnectome map and number of voxels with disconnection probability >50%) were strongly correlated with lesion size (posterior mean = 0.74, BF >150 and median posterior *δ* = 0.68, BF >150, respectively). Global disconnectivity (mean value in disconnectome map) was not correlated with FSS (median posterior *δ* = 0.03) or PHQ (median posterior *δ* = 0.03). Bayesian correlations (using default priors) provided moderate evidence (BF = 0.26) for these null effects, relative to H1 (positive associations between disconnectivity measures and FSS/PHQ). This indication of a null effect was mirrored in correlations between the alternative measure of disconnectivity (number of voxels with disconnection probability >50%) and FSS (median posterior *δ* = 0.05, BF = 0.29), and PHQ (median posterior *δ* = 0.02, BF = 0.26). For transparency, Bayes factors of the main correlations estimated on different priors are reported in Supplementary Table 1, while Supplementary Table 2 reports the correlations after removing the most extreme outlier in terms of lesion size.

### Associations between clinical stroke-related characteristics and FSS

Linear models (lmBF) corrected for age and depressive symptoms did not provide evidence for associations between FSS scores and lesion location (brainstem/cerebellum, left or right hemisphere or both hemispheres), lesion volume, months since stroke or TOAST stroke classification (see Supplementary Table 3 for model comparisons and associated Bayes factors). No stroke related variable, including global disconnectivity, demonstrated Bayes factors >1, indicating low predictive value for all. Specifically, all extended models with stroke lesion variables displayed Bayes factors below 0.33 when compared to the simpler null model, suggesting moderate evidence of no effect of stroke lesion characteristics.

## Discussion

Fatigue following stroke is common and represents a significant clinical burden. Stroke sequelae reflect both cell death at the site of the lesion, as well as structural and functional alterations in extended brain networks. Brain network dysfunction, directly or indirectly related to the stroke lesion, is a putative mechanism underlying PSF pathophysiology. Previous studies have primarily assessed lesion characteristics such as volume or location, and the added explanatory value of probing the extended brain network connections with the lesion has been unclear. To this end, hypothesizing that network disruptions in cortico-striatal networks would be associated with higher levels of fatigue, we calculated disconnectivity maps for 84 patients in the chronic phase and used permutation testing to evaluate the association between PSF symptoms and regional network disconnection.

Permutation testing revealed no significant associations between symptoms of fatigue and disconnectome maps or binarized lesion maps. We found no support for our hypothesis that a disconnectivity approach by disconnectome maps would add predictive value of fatigue beyond conventional lesion analyses. In line with this, Bayes factor estimations on correlations between disconnectivity summary measures and FSS score provided moderate evidence for the null hypothesis (no association) relative to the alternative hypothesis (association between fatigue and disconnectivity). However, results are not decisive, and alternative explanations of the absent effects must be considered.

The lack of added predictive value from the disconnectivity measures when compared to more traditional lesion characteristics is in general agreement with recent studies (Hope, Leff, & Price, 2018; Salvalaggio et al., 2020) reporting similar predictive value for models including (dis)connectivity measures compared to models with lesion information only. The lack of robust associations between disconnectome maps and the clinical measures has several likely explanations. It has been suggested that the information provided in the disconnectome maps is largely embedded in the binarized lesion masks (Hope et al., 2018; Salvalaggio et al., 2020), implying that the two representations of lesion related pathology convey overlapping variance. Indeed, the correlation between lesion volume and global disconnectivity, operationalized as the average voxel value across the disconnectome map or the number of voxels with probability of disconnection >50%, was relatively strong, intuitively supporting that larger lesions project to a larger proportion of the brain.

Alternatively, it may be that disconnectome maps and lesion masks convey similar information primarily when the sample is large and lesion diversity sufficiently high to capture the variance embedded in the disconnectome maps (Griffis et al., 2019). This could imply that in many real-life situations where large samples are not always realistic/feasible, such as in clinical stroke populations, disconnectome maps may provide relevant, complimentary and unique information. In agreement with this, a recent study revealed that structural disconnectivity maps explained a larger proportion of the variance in core functional connectivity disruptions than did focal lesions, and displayed significant correlations with behavior (Griffis et al., 2019), thus facilitating the understanding of individual differences in outcome. Moreover, a study by Kuceyeski et al. (2016) reported higher accuracy in cognitive and mobility prediction for models including disconnection metrics than models based on lesion volume.

A key assumption underlying our analyses is that the indirectly calculated disconnectome maps provide a realistic estimate of structural network disconnection and that these disconnections have functional effects. As depicted in Figure 3, the degree of estimated tract disconnection can be extensive, even for smaller lesions. While such lesion to brain network mapping supports the notion that lesions in hub-like regions project to and implicates an extended set of brain regions and networks (Colizza, Flammini, Serrano, & Vespignani, 2006; Van Den Heuvel & Sporns, 2011), the tractography process used to build the normative training set has several inherent limitations and errors can be introduced in any stage of the tracking process (Schilling et al., 2019). Noise and artefacts in the image acquisition, difficulties establishing fiber orientation (Jeurissen, Descoteaux, Mori, & Leemans, 2019) and choices regarding the tracking algorithm and parameters such as stop criteria and curvature threshold (Knösche, Anwander, Liptrot, & Dyrby, 2015; Schilling et al., 2019), are among the commonly recognized pitfalls. Consequently, the reconstructed pathways based on diffusion tractography may not necessarily reflect true structural connections, and to which degree disconnectome maps reflect biological disconnections is still debated (Salvalaggio et al., 2020), warranting caution when interpreting tractography results without supporting converging evidence (Jeurissen et al., 2019). These limitations are not specific for the currently employed algorithm, and further work is needed to overcome general limitation of biological accuracy and validity of diffusion based tractography.

The added value of disconnectome maps in brain-behavior mapping also depends on the reliability, validity and functional neuroanatomy of the included clinical and behavioral measures. For example, primary motor dysfunctions, which may require simpler operationalizations and measurements than more complex cognitive symptoms, are more strongly associated with focal damage, while other behaviors, like verbal associative memory, may be more strongly predicted by extended network function (Griffis et al., 2019; Siegel et al., 2016). The lack of significant associations between brain characteristics and behavioral measures in the current study may therefore be partly related to the properties and measurement of PSF. Although fatigue is painfully tangible for the individual patient, it is it is unspecific and difficult to operationalize, and the lack of “gold standard” measures of subjective fatigue has been characterized as one of the major obstacles to PSF research (Nadarajah & Goh, 2015). In the current study, we applied the FSS as a general measure of fatigue interference and severity. As FSS is the most widely used fatigue measure in stroke research (Cumming et al., 2016), reporting FSS scores facilitates communication and synthesizing of results across studies. Still, FSS constitutes a rather coarse measure of a complex phenomenon, and does not provide information on other relevant aspects of PSF such as diurnal fluctuations and fatigue subtypes. It is conceivable that more finely grained measures of i.e. fatigue subtypes could reveal associations not detected by the FSS.

Mimicking the results from the disconnectome approach, linear regressions testing for associations between FSS and lesion characteristics (volume and location) revealed no significant associations. This is in agreement with several previous studies (Choi-Kwon et al., 2005; Ingles et al., 1999; Mead et al., 2011). Still, the literature is inconclusive, and some suggest significant associations between PSF and lesion characteristics (Snaphaan et al., 2011; Wai Kwong Tang et al., 2014; Wai Kwong Tang et al., 2010). The inconsistency between studies may be attributable to differences in how lesion site is defined and reported, as well as time since stroke and clinical characteristics and severity of the sample. Studies that do not report significant associations between fatigue and lesion characteristics tend to define lesion location broadly (Wu, Mead, Macleod, & Chalder, 2015), such as posterior/anterior circulation or left/right hemisphere, while studies reporting significant associations often apply a more detailed account of lesion site (e.g. which specific structures are affected) and are conducted within the first few months after stroke. The temporal aspect may be of particular importance, considering that the character of stroke sequelae and associated brain correlates change over time through processes of recovery and compensation (Fornito, Zalesky, & Breakspear, 2015; Fox, 2018). In the present study, fatigue was measured on average 22 months post stroke. The absence of identified stroke lesion effects may thus suggest that lesion characteristics play a less critical role in the chronic phase (Aarnes, Stubberud, & Lerdal, 2020).

In addition to the general limitations related to the interpretation of imaging-based measures of brain connectivity listed above, the results should be interpreted in light of the following caveats. First, the recruited patients suffered mild strokes and were drawn from a highly functioning part of the stroke population, as the extent and type of assessments prevented the more disabled patients from participating (e.g. severe aphasia, paralysis, severe neglect). This limits generalizability of results, and we cannot exclude the possibility that including more severely fatigued patients would reveal associations not detected in the current study. However, even in this sample of fairly high functioning stroke patients, levels of fatigue were significantly higher than in the healthy control group, and comparable to fatigue levels reported in other samples of chronic stroke patients (Choi-Kwon et al., 2005; Valko, Bassetti, Bloch, Held, & Baumann, 2008). Moreover, fatigue correlated strongly with depression, in line with previous reports (Choi-Kwon et al., 2005; van de Port, Kwakkel, Bruin, & Lindeman, 2007), intuitively indicating that FSS scores reflect a relevant clinical phenotype.

Second, VSLM analyses are inherently contingent on and restricted by the variability of the patients’ lesion locations, as a lesion site cannot be identified as important for a symptom if it is not represented in the sample. With regards to the current sample, the lack of whole brain representation limits the spatial scope of the analyses, where i.e. right hemispheric strokes were more densely sampled than left hemispheric strokes, and prefrontal cortex was marginally affected. This lack of full or random sampling of the brain represents a common caveat in stroke research, because stroke lesions are not randomly or evenly dispersed throughout the brain, but are dependent on vascular organization and architecture and tend to occur in proximity to major arteries (Rorden, Karnath, & Bonilha, 2007; Zhao, Halai, & Lambon Ralph, 2020). In line with this, degree of voxel-wise lesion overlap between patients in the current sample was low, and although a sample size of 84 is comparable with common practice in MRI studies targeting stroke (Nickel & Thomalla, 2017; Nott et al., 2019; Sihvonen, Ripollés, Rodríguez-Fornells, Soinila, & Särkämö, 2017), further studies with even larger samples are needed.

Low power due to small sample sizes is common in neuroscience (Button et al., 2013), and might be especially pressing in stroke imaging research where inter-patient variability in lesions and symptoms is high, and large datasets are logistically and financially demanding to collect (Liew et al., 2020; Price, Hope, & Seghier, 2017). With reference to this fundamental constraint, the best hope for future stroke neuroimaging studies may lie in large-scale data-sharing initiatives such as the ENIGMA Stroke Recovery database (Liew et al., 2020), where pooled and synthesized data from individual studies facilitates conduction of well powered studies on large and diverse samples.

However, in smaller samples with low lesion overlap, targeting disconnectivity through disconnectome maps may be particularly relevant, because such measures reveal common disruptions across spatially dispersed lesions (Griffis et al., 2019), resulting in a higher degree of disconnectome overlap compared to lesion overlap.

In conclusion, the current study represents a novel approach to assess the neural correlates of PSF in chronic stroke patients. By indirectly estimating structural network disconnections caused by the stroke lesions, we arrived at individual disconnectome maps capturing distal effects of focal damage. The results did not provide evidence that a disconnectome based approach demonstrates higher sensitivity to PSF than a VLSM approach. Nor did the results support the notion that lesions to particular regions or disconnections to specific networks contribute to PSF in the chronic phase. However, methodological considerations regarding statistical power, lesion coverage- and overlap warrants caution when interpreting results.

## Supporting information

supplementary Materialt

This study was carried out in accordance with The Code of Ethics of the World Medical Association (Declaration of Helsinki)

## Acknowledgements

This study was supported by the Norwegian ExtraFoundation for Health and Rehabilitation (2015/FO5146), the Research Council of Norway (249795, 262372), the South □ Eastern Norway Regional Health Authority (2014097, 2015044, 2015073, 2018037), the European Research Council under the European Union’s Horizon 2020 research and Innovation program (ERC StG Grant 802998), and the Department of Psychology, University of Oslo.

## Author contributions

KMU conceived the study, collected the data, analyzed the data and wrote the manuscript. KK and GR collected the data and were involved in data quality control and pre-processing of MRI images, DA created main figures and was involved in critical revision of the paper, ESD and JM collected data and were involved in data quality control and performed the lesion segmentations, MTS estimated the disconnectome maps and critically revised the paper, AMS were involved in data collection, quality control and contributed to drafting the paper, HIH, ST, AE and JEN were involved in interpreting of results and critical revision of the paper, LTW was involved in conception of the study, data analysis, interpretation of results and critical revision of manuscript.

## References

Adams Jr, H. P., Bendixen, B. H., Kappelle, L. J., Biller, J., Love, B. B., Gordon, D. L., & Marsh 3rd, E. E. (1993). Classification of subtype of acute ischemic stroke. Definitions for use in a multicenter clinical trial. TOAST. Trial of Org 10172 in Acute Stroke Treatment. Stroke, 24(1), 35–41.

Association, U. S. (2020). Retrieved from https://www.stroke.org.uk/effects-of-stroke/tiredness-and-fatigue#What%20is%20post-stroke%20fatigue?

Avants, B. B., Tustison, N. J., Song, G., Cook, P. A., Klein, A., & Gee, J. C. (2011). A reproducible evaluation of ANTs similarity metric performance in brain image registration. NeuroImage, 54(3), 2033–2044.

Bartolomeo, P., & de Schotten, M. T. (2016). Let thy left brain know what thy right brain doeth: inter-hemispheric compensation of functional deficits after brain damage. Neuropsychologia, 93, 407–412.

Bates, E., Wilson, S. M., Saygin, A. P., Dick, F., Sereno, M. I., Knight, R. T., & Dronkers, N. F. (2003). Voxel-based lesion–symptom mapping. Nature neuroscience, 6(5), 448–450.

Bonilha, L., Nesland, T., Rorden, C., Fillmore, P., Ratnayake, R. P., & Fridriksson, J. (2014). Mapping remote subcortical ramifications of injury after ischemic strokes. Behavioural neurology, 2014.

Brain Connectivity Behaviour Toolkit (BCBtoolkit). Retrieved from http://toolkit.bcblab.com

Button, K. S., Ioannidis, J. P. A., Mokrysz, C., Nosek, B. A., Flint, J., Robinson, E. S. J., & Munafò, M. R. (2013). Power failure: why small sample size undermines the reliability of neuroscience. Nature Reviews Neuroscience, 14(5), 365–376.

Carrera, E., & Tononi, G. (2014). Diaschisis: past, present, future. Brain, 137(9), 2408–2422.

Chaudhuri, A., & Behan, P. O. (2000). Fatigue and basal ganglia. Journal of the neurological sciences, 179(1-2), 34–42.

Chaudhuri, A., & Behan, P. O. (2004). Fatigue in neurological disorders. The lancet, 363(9413), 978–988.

Choi-Kwon, S., Han, S. W., Kwon, S. U., & Kim, J. S. (2005). Poststroke fatigue: characteristics and related factors. Cerebrovascular Diseases, 19(2), 84–90.

Colizza, V., Flammini, A., Serrano, M. A., & Vespignani, A. (2006). Detecting rich-club ordering in complex networks. Nature physics, 2(2), 110–115.

Cotovio, G., Talmasov, D., Barahona-Corrêa, J. B., Hsu, J., Senova, S., Ribeiro, R., Rost, N. (2020). Mapping mania symptoms based on focal brain damage. The Journal of Clinical Investigation, 130(10).

Cowan, W. M. (1970). Anterograde and retrograde transneuronal degeneration in the central and peripheral nervous system. In Contemporary research methods in neuroanatomy (pp. 217–251): Springer.

Cumming, T. B., Packer, M., Kramer, S. F., & English, C. (2016). The prevalence of fatigue after stroke: a systematic review and meta-analysis. International Journal of stroke, 11(9), 968–977.

Cumming, T. B., Yeo, A. B., Marquez, J., Churilov, L., Annoni, J.-M., Badaru, U., Lerdal, A. (2018). Investigating post-stroke fatigue: An individual participant data meta-analysis. Journal of psychosomatic research, 113, 107–112.

Darby, R. R., Joutsa, J., & Fox, M. D. (2019). Network localization of heterogeneous neuroimaging findings. Brain, 142(1), 70–79.

de Bruijn, M. A. A. M., Synhaeve, N. E., van Rijsbergen, M. W. A., de Leeuw, F.-E., Mark, R. E., Jansen, B. P. W., & de Kort, P. L. M. (2015). Quality of life after young ischemic stroke of mild severity is mainly influenced by psychological factors. Journal of Stroke and Cerebrovascular Diseases, 24(10), 2183–2188.

De Doncker, W., Dantzer, R., Ormstad, H., & Kuppuswamy, A. (2018). Mechanisms of poststroke fatigue. Journal of Neurology, Neurosurgery & Psychiatry, 89(3), 287–293.

de Groot, M. H., Phillips, S. J., & Eskes, G. A. (2003). Fatigue associated with stroke and other neurologic conditions: implications for stroke rehabilitation. Archives of physical medicine and rehabilitation, 84(11), 1714–1720.

Douven, E., Köhler, S., Schievink, S. H. J., van Oostenbrugge, R. J., Staals, J., Verhey, F. R. J., & Aalten, P. (2017). Temporal associations between fatigue, depression, and apathy after stroke: results of the cognition and affect after stroke, a prospective evaluation of risks study. Cerebrovascular Diseases, 44(5-6), 330–337.

Dørum, E. S., Kaufmann, T., Alnæs, D., Richard, G., Kolskår, K. K., Engvig, A., Nordvik, J. E. (2020). Functional brain network modeling in sub-acute stroke patients and healthy controls during rest and continuous attentive tracking. Heliyon, 6(9), e04854.

Ferguson, M. A., Lim, C., Cooke, D., Darby, R. R., Wu, O., Rost, N. S., Fox, M. D. (2019). A human memory circuit derived from brain lesions causing amnesia. Nature communications, 10(1), 1–9.

Fornito, A., Zalesky, A., & Breakspear, M. (2015). The connectomics of brain disorders. Nature Reviews Neuroscience, 16(3), 159–172.

Foulon, C., Cerliani, L., Kinkingnehun, S., Levy, R., Rosso, C., Urbanski, M., Thiebaut de Schotten, M. (2018). Advanced lesion symptom mapping analyses and implementation as BCBtoolkit. GigaScience, 7(3), giy004.

Fox, M. D. (2018). Mapping symptoms to brain networks with the human connectome. New England Journal of Medicine, 379(23), 2237–2245.

Geschwind, N. (1974). Disconnexion syndromes in animals and man. In Selected papers on language and the brain (pp. 105–236): Springer.

Glader, E.-L., Stegmayr, B., & Asplund, K. (2002). Poststroke fatigue: a 2-year follow-up study of stroke patients in Sweden. Stroke, 33(5), 1327–1333.

Griffis, J. C., Metcalf, N. V., Corbetta, M., & Shulman, G. L. (2019). Structural disconnections explain brain network dysfunction after stroke. Cell reports, 28(10), 2527–2540.

Hope, T. M. H., Leff, A. P., & Price, C. J. (2018). Predicting language outcomes after stroke: Is structural disconnection a useful predictor? NeuroImage: Clinical, 19, 22–29.

Høgestøl, E. A., Nygaard, G. O., Alnæs, D., Beyer, M. K., Westlye, L. T., & Harbo, H. F. (2019). Symptoms of fatigue and depression is reflected in altered default mode network connectivity in multiple sclerosis. PLoS One, 14(4), e0210375.

Ingles, J. L., Eskes, G. A., & Phillips, S. J. (1999). Fatigue after stroke. Archives of physical medicine and rehabilitation, 80(2), 173–178.

Jeurissen, B., Descoteaux, M., Mori, S., & Leemans, A. (2019). Diffusion MRI fiber tractography of the brain. NMR in Biomedicine, 32(4), e3785.

Keysers, C., Gazzola, V., & Wagenmakers, E.-J. (2020). Using Bayes factor hypothesis testing in neuroscience to establish evidence of absence. Nature neuroscience, 23(7), 788–799.

Klein, A., Andersson, J., Ardekani, B. A., Ashburner, J., Avants, B., Chiang, M.-C., Hellier, P. (2009). Evaluation of 14 nonlinear deformation algorithms applied to human brain MRI registration. NeuroImage, 46(3), 786–802.

Knösche, T. R., Anwander, A., Liptrot, M., & Dyrby, T. B. (2015). Validation of tractography: comparison with manganese tracing. Human brain mapping, 36(10), 4116–4134.

Kolskaar, K. K., Richard, G., Alnaes, D., Dørum, E. S., Sanders, A.-M., Ulrichsen, K. M., Westlye, L. T. (2020). Reliability, sensitivity and predictive value of fMRI during multiple object tracking as a marker of cognitive training in stroke survivors. bioRxiv, 603985.

Kroenke, K., Spitzer, R. L., & Williams, J. B. W. (2001). The PHQ-9: validity of a brief depression severity measure. Journal of general internal medicine, 16(9), 606–613.

Krupp, L. B., LaRocca, N. G., Muir-Nash, J., & Steinberg, A. D. (1989). The fatigue severity scale: application to patients with multiple sclerosis and systemic lupus erythematosus. Archives of neurology, 46(10), 1121–1123.

Kuceyeski, A., Navi, B. B., Kamel, H., Raj, A., Relkin, N., Toglia, J., O’Dell, M. (2016). Structural connectome disruption at baseline predicts 6lJmonths postlJstroke outcome. Human brain mapping, 37(7), 2587–2601.

Kunze, A., Zierath, D., Drogomiretskiy, O., & Becker, K. (2014). Strain differences in fatigue and depression after experimental stroke. Translational stroke research, 5(5), 604–611.

Lerdal, A., Bakken, L. N., Rasmussen, E. F., Beiermann, C., Ryen, S., Pynten, S., Finset, A. (2011). Physical impairment, depressive symptoms and pre-stroke fatigue are related to fatigue in the acute phase after stroke. Disability and rehabilitation, 33(4), 334–342.

Liew, S. L., Zavaliangos-Petropulu, A., Jahanshad, N., Lang, C. E., Hayward, K. S., Lohse, K. R., . . Bhattacharya, A. K. (2020). The ENIGMA Stroke Recovery Working Group: Big data neuroimaging to study brain–behavior relationships after stroke. Human brain mapping.

Lyden, P., Raman, R., Liu, L., Emr, M., Warren, M., & Marler, J. (2009). National Institutes of Health Stroke Scale certification is reliable across multiple venues. Stroke, 40(7), 2507–2511.

Mead, G. E., Graham, C., Dorman, P., Bruins, S. K., Lewis, S. C., Dennis, M. S., IST, U. K. C. o. (2011). Fatigue after stroke: baseline predictors and influence on survival. Analysis of data from UK patients recruited in the International Stroke Trial. PLoS One, 6(3).

Morey, R. D., Rouder, J. N., Jamil, T., & Morey, M. R. D. (2015). Package ‘bayesfactor’. *URLh http://cran/r-projectorg/web/packages/BayesFactor/BayesFactor pdf i (accessed 1006 15)*.

Mutai, H., Furukawa, T., Houri, A., Suzuki, A., & Hanihara, T. (2017). Factors associated with multidimensional aspect of post-stroke fatigue in acute stroke period. Asian journal of psychiatry, 26, 1–5.

Nadarajah, M., & Goh, H.-T. (2015). Post-stroke fatigue: a review on prevalence, correlates, measurement, and management. Topics in stroke rehabilitation, 22(3), 208–220.

Naess, H., Lunde, L., Brogger, J., & Waje-Andreassen, U. (2012). Fatigue among stroke patients on long-term follow-up. The Bergen Stroke Study. Journal of the neurological sciences, 312(1-2), 138–141.

Naess, H., Waje-Andreassen, U., Thomassen, L., Nyland, H., & Myhr, K.-M. (2006). Health-related quality of life among young adults with ischemic stroke on long-term follow-up. Stroke, 37(5), 1232–1236.

Nasreddine, Z. S. (2020). Retrieved from https://www.mocatest.org

Nasreddine, Z. S., Phillips, N. A., Bédirian, V., Charbonneau, S., Whitehead, V., Collin, I., Chertkow, H. (2005). The Montreal Cognitive Assessment, MoCA: a brief screening tool for mild cognitive impairment. Journal of the American Geriatrics Society, 53(4), 695–699.

Nickel, A., & Thomalla, G. (2017). Post-stroke depression: impact of lesion location and methodological limitations—a topical review. Frontiers in Neurology, 8, 498.

Nordin, L. E., Möller, M. C., Julin, P., Bartfai, A., Hashim, F., & Li, T.-Q. (2016). Post mTBI fatigue is associated with abnormal brain functional connectivity. Scientific reports, 6, 21183.

Nott, Z., Horne, K., Prangley, T., Adams, A. G., Henry, J. D., & Molenberghs, P. (2019). Structural and functional brain correlates of theory of mind impairment post-stroke. Cortex, 121, 427–442.

Ponchel, A., Bombois, S., Bordet, R., & Hénon, H. (2015). Factors associated with poststroke fatigue: a systematic review. Stroke research and treatment, 2015.

Price, C. J., Hope, T. M., & Seghier, M. L. (2017). Ten problems and solutions when predicting individual outcome from lesion site after stroke. NeuroImage, 145, 200–208.

R Core Team. (2017). R: A language and environment for statistical computing. R Foundation for Statistical Computing. Vienna, Austria. Retrieved from URL https://www.R-project.org/.

Rehme, A. K., & Grefkes, C. (2013). Cerebral network disorders after stroke: evidence from imaging-based connectivity analyses of active and resting brain states in humans. The Journal of physiology, 591(1), 17–31.

Richard, G., Kolskår, K., Sanders, A.-M., Kaufmann, T., Petersen, A., Doan, N. T., Dørum, E. S. (2018). Assessing distinct patterns of cognitive aging using tissue-specific brain age prediction based on diffusion tensor imaging and brain morphometry. PeerJ, 6, e5908.

Rorden, C., Karnath, H.-O., & Bonilha, L. (2007). Improving lesion-symptom mapping. Journal of cognitive neuroscience, 19(7), 1081–1088.

Salvalaggio, A., De Filippo de Grazia, M., Zorzi, M., Thiebaut de Schotten, M., & Corbetta, M. (2020). Post-stroke deficit prediction from lesion and indirect structural and functional disconnection. Brain, 143(7), 2173–2188.

Schepers, V. P., Visser-Meily, A. M., Ketelaar, M., & Lindeman, E. (2006). Poststroke fatigue: course and its relation to personal and stroke-related factors. Archives of physical medicine and rehabilitation, 87(2), 184–188.

Schilling, K. G., Nath, V., Hansen, C., Parvathaneni, P., Blaber, J., Gao, Y., Ocampo-Pineda, M. (2019). Limits to anatomical accuracy of diffusion tractography using modern approaches. NeuroImage, 185, 1–11.

Siegel, J. S., Ramsey, L. E., Snyder, A. Z., Metcalf, N. V., Chacko, R. V., Weinberger, K., Corbetta, M. (2016). Disruptions of network connectivity predict impairment in multiple behavioral domains after stroke. Proceedings of the National Academy of Sciences, 113(30), E4367–E4376.

Sihvonen, A. J., Ripollés, P., Rodríguez-Fornells, A., Soinila, S., & Särkämö, T. (2017). Revisiting the neural basis of acquired amusia: lesion patterns and structural changes underlying amusia recovery. Frontiers in neuroscience, 11, 426.

Smith, S. M., & Nichols, T. E. (2009). Threshold-free cluster enhancement: addressing problems of smoothing, threshold dependence and localisation in cluster inference. NeuroImage, 44(1), 83–98.

Snaphaan, L., Van der Werf, S., & de Leeuw, F. E. (2011). Time course and risk factors of post-stroke fatigue: a prospective cohort study. European Journal of Neurology, 18(4), 611–617.

Spitzer, R. L., Kroenke, K., Williams, J. B. W., & Study, G. (1999). Validation and utility of a self-report version of PRIME-MD: the PHQ primary care study. Jama, 282(18), 1737–1744.

Stuart, E. A., King, G., Imai, K., & Ho, D. (2011). MatchIt: nonparametric preprocessing for parametric causal inference. Journal of statistical software.

Tang, W. K., Chen, Y. K., Liang, H. J., Chu, W. C. W., Mok, V. C. T., Ungvari, G. S., & Wong, K. S. (2014). Subcortical white matter infarcts predict 1-year outcome of fatigue in stroke. BMC neurology, 14(1), 234.

Tang, W. K., Chen, Y. K., Mok, V., Chu, W. C. W., Ungvari, G. S., Ahuja, A. T., & Wong, K. S. (2010). Acute basal ganglia infarcts in poststroke fatigue: an MRI study. Journal of neurology, 257(2), 178–182.

Tang, W. K., Liang, H. J., Chen, Y. K., Chu, W. C. W., Abrigo, J., Mok, V. C. T., Wong, K. S. (2013). Poststroke fatigue is associated with caudate infarcts. Journal of the neurological sciences, 324(1-2), 131–135.

Thiebaut de Schotten, M., Foulon, C., & Nachev, P. (2020). Brain disconnections link structural connectivity with function and behaviour. Nature communications, 11(1), 5094. doi:10.1038/s41467-020-18920-9

Thomas, K., Gamlin, C., de Simoni, A., Mullis, R., & Mant, J. (2019). How is poststroke fatigue understood by stroke survivors and carers? A thematic analysis of an online discussion forum. BMJ open, 9(7), e028958.

Valko, P. O., Bassetti, C. L., Bloch, K. E., Held, U., & Baumann, C. R. (2008). Validation of the fatigue severity scale in a Swiss cohort. Sleep, 31(11), 1601–1607.

van de Port, I. G. L., Kwakkel, G., Bruin, M., & Lindeman, E. (2007). Determinants of depression in chronic stroke: a prospective cohort study. Disability and rehabilitation, 29(5), 353–358.

Van Den Heuvel, M. P., & Sporns, O. (2011). Rich-club organization of the human connectome. Journal of Neuroscience, 31(44), 15775–15786.

von Monakow, C. (1914). Die Lokalisation im Grosshirn und der Abbau der Funktion durch kortikale Herde: JF Bergmann.

Wang, R., Benner, T., Sorensen, A. G., & Wedeen, V. J. (2007). Diffusion toolkit: a software package for diffusion imaging data processing and tractography.

Whitehead, L. (2009). The measurement of fatigue in chronic illness: a systematic review of unidimensional and multidimensional fatigue measures. Journal of pain and symptom management, 37(1), 107–128.

Winkler, A. M., Ridgway, G. R., Webster, M. A., Smith, S. M., & Nichols, T. E. (2014). Permutation inference for the general linear model. NeuroImage, 92, 381–397.

Winward, C., Sackley, C., Metha, Z., & Rothwell, P. M. (2009). A population-based study of the prevalence of fatigue after transient ischemic attack and minor stroke. Stroke, 40(3), 757–761.

Wu, S., Barugh, A., Macleod, M., & Mead, G. (2014). Psychological associations of poststroke fatigue: a systematic review and meta-analysis. Stroke, 45(6), 1778–1783.

Wu, S., Mead, G., Macleod, M., & Chalder, T. (2015). Model of understanding fatigue after stroke. Stroke, 46(3), 893–898.

Zhao, Y., Halai, A. D., & Lambon Ralph, M. A. (2020). Evaluating the granularity and statistical structure of lesions and behaviour in post-stroke aphasia. Brain Communications, 2(2), fcaa062.

Aarnes, R., Stubberud, J., & Lerdal, A. (2019). A literature review of factors associated with fatigue after stroke and a proposal for a framework for clinical utility. Neuropsychological rehabilitation, 1–28.

Aarnes, R., Stubberud, J., & Lerdal, A. (2020). A literature review of factors associated with fatigue after stroke and a proposal for a framework for clinical utility. Neuropsychological rehabilitation, 30(8), 1449–1476.

