## supplementary Materialt for "Brain disconnectivity mapping of post-stroke fatigue"

**Supplementary material**

| Table 1 | | | |
| --- | --- | --- | --- |
| *Varying priors on disconnectivity models* | Prior | BF | *δ*  & 95% CI |
| corrBF (FSS, mean_disconnectivity) | MediumNarrow | 0.34 | 0.02 [-0.17 – 0.23] |
| corrBF (FSS, mean_disconnectivity)* | **DefaultMedium** | **0.26** | 0.03 [-0.18 – 0.24] |
| corrBF (FSS, mean_disconnectivity) | Wide | 0.20 | 0.03 [-0.18 – 0.24] |
| corrBF(FSS, n voxels > 50% disconnect) | MediumNarrow | 0.38 | 0.05 [-0.15 – 0.26] |
| corrBF(FSS, n voxels > 50% disconnect)* | **DefaultMedium** | **0.29** | 0.06 [-0.15 – 0.27] |
| corrBF(FSS, n voxels > 50% disconnect) | Wide | 0.22 | 0.05 [-0.15 – 0.27] |
| *Models reported in manuscript |  |  |  |

| Table 2  *Main models with outlier excluded* | BF | *δ*  & 95% CI |
| --- | --- | --- |
| corrBF (FSS, mean_disconnectivity) | 0.32 | 0.07 [-0.13 – 0.29] |
| corrBF (PHQ, mean_disconnectivity) | 0.26 | 0.01 [-0.19 – 0.22] |
| corrBF (FSS, n voxels > 50% disconnect) | 0.39 | 0.09 [-0.11 – 0.30] |
| corrBF(PHQ, n voxels > 50% disconnect) | 0.29 | 0.05 [-0.16 – 0.26] |
| corrBF(FSS, number of voxels lesioned) | 0.28 | 0.05 [-0.16 – 0.25] |
| corrBF(PHQ, number of voxels in lesion) | 0.29 | 0.05 [-0.16 – 0.26] |

| Table 3 |  |
| --- | --- |
| **Model comparison by Bayes Factor**  *One variable added in each model and compared against null model* | Bayes Factor |
| **“Null model FSS”: lmBF(FSS ~ Age + PHQ) / intercept only** | >150 ±0% |
| Null + sex / null | 0.29 ±0.28% |
| Null + months since stroke / null | 0.17 ±0% |
| Null + TOAST / null | 0.16 ±0.42% |
| Null + lesion location / null | 0.19 ±0.38% |
| Null + lesion size / null | 0.16 ±0% |
| Null + mean disconnectivity / null | 0.19 ±0% |
| Null + number of voxels disconnectivity probability >50% / null | 0.19 ±0% |

No models (except “null model”) display Bayes factors >1, indicating that variables have low predictive value and that no extended models are preferred over the null model.
